# GBSmode: a pipeline for haplotype-aware analysis of genotyping-by-sequencing data

**DOI:** 10.1101/2021.09.20.461130

**Authors:** Steven Yates, Bruno Studer

## Abstract

2

**Background:** Genotyping-by-sequencing (GBS) has revolutionised molecular genetic analysis. It enables simultaneous genotyping of thousands of DNA markers in the genome of any species. In contrast to whole-genome shotgun sequencing, GBS exploits a restriction enzyme to reduce genome complexity and directs the sequencing to begin at fixed digestion sites. However, currently used tools for the analysis of GBS data, such as SAMtools, often neglect the fundamental technical differences between GBS and shotgun sequencing.

**Results:** Here we present GBSmode, a dedicated pipeline to call DNA sequence variants using whole-read information from GBS data. It removes false positives by incorporating biological features such as the ploidy level and the number of possible alleles in the population under investigation. Comparison of GBSmode with SAMtools in an F2 population of rice (*Oryza sativa* L.) showed both identified a similar number of polymorphisms (13,449 and 14,445, respectively) with a high overlap (8,143). However, differences were found in the number of read misalignments (8.0% and 14.3% for GBSmode and SAMtools, respectively) and genotyping errors (5.0% and 8.3% for GBSmode and SAMtools, respectively). Further tests in a bi-parental F1 population of cassava (*Manihot esculenta* Crantz) showed GBSmode found 31,489 polymorphic loci, whereas the number was higher with SAMtools (43,860). However, this difference was mainly attributable to GBSmode rejecting 11,695 loci that were biologically not possible.

**Conclusions:** This study shows that GBSmode is a versatile tool for the analysis of GBS data. Moreover, GBSmode was able to reduce genotyping errors arising from read misalignments by combining haplotype data with biological information. Whilst other tools may find more markers, GBSmode is designed for accuracy.

## 4 Background

Genotyping-by-sequencing (GBS) is a cost-effective method for genotyping thousands of markers simultaneously in large populations of human, plant and animal species. It combines restriction enzyme-based genome complexity reduction with modern sequencing technologies (Scheben *et al*., 2017). The GBS method reported by Elshire *et al*., (2011) is a four-step protocol (digestion, ligation, amplification and sequencing) that efficiently enables multiplexing of hundreds of samples. GBS is mainly used to identify single nucleotide polymorphisms (SNPs) for diversity analyses, linkage mapping and marker-trait association studies (Scheben *et al*., 2017). Compared with array-based genotyping methods, GBS is competitive in price and advantageous in orphan crop species (Rasheed *et al*., 2017). Despite high missing value rates and a higher computational demand required for data analysis, GBS has evolved as the tool of choice for many applications.

The core principle of GBS is the use of a restriction enzyme to fragment the genome, followed by the enrichment of low molecular weight fragments of around 250 bp using PCR amplification (Elshire *et al*., 2011). GBS is often termed a ‘reduced representation/complexity library’ which is attributable to GBS’s repeated sampling of the same segments of the genome (Scheben *et al*., 2017) among individuals. From a technical perspective, GBS directs the sequencing to commence at specific sites, enabling site-directed sequencing. This is different to shotgun sequencing, used for whole genome sequencing (WGS, Ekblom & Wolf, 2014) and transcriptome sequencing (Mortazavi *et al*., 2008). As the name implies, shotgun sequencing uses randomly fragmented DNA as a starting point, meaning that the library preparation is fundamentally different compared to GBS.

Aligning short DNA reads to a genome sequence is error prone, especially when the short read comes from a repetitive genome region. Shotgun sequencing can tackle the read alignment ambiguity by aligning reads across a region, to find enough differences to unambiguously place reads. This means many reads (coverage) are needed to build a coherent picture of any loci, from reads that surround the loci. From these assumptions, a majority rule applies and the most frequently occurring base is considered the correct one. Yet GBS data differs fundamentally in this regard because DNA is digested at specific recognition sites, meaning that reads are uniformly stacked on a locus.

Another important but often neglected aspect is that GBS reads may contain multiple polymorphisms. Standard tools for the analysis of next generation sequencing (NGS) data identify variants independently from one base pair to another and omits neighbouring information. But if a read contains multiple polymorphisms, standard tools, that use ‘pileups’, lose this information. However, making use of polymorphisms found together on the same NGS read can increase both the precision of variant calling and the number of alleles at a site. For the former, multiple polymorphisms can mitigate sequencing errors and increase the detection confidence of real polymorphisms. For the latter, multiple polymorphisms can be used to form haplotypes that better represent multi-allelic populations, better than bi-morphic polymorphisms. This is useful for outcrossing species, where a high frequency of polymorphisms is often found. For example, in cassava (*Manihot esculenta* Crantz), 26 million SNPs were identified in 241 accessions by WGS (Ramu *et al*., 2017).

Our aim was to develop a dedicated workflow to analyse GBS data which incorporates i) the information of the digestion site, ii) the information of polymorphisms found together in the same read to identify haplotypes and iii) the biological information of the population under investigation to filter putative haplotypes and increase accuracy. GBSmode, implementing these aims in an open source R-pipeline, was compared with SAMtools: on two publicly available datasets.

## 5 Implementation

### 5.1 Datasets

The first dataset used to compare the performance of GBSmode with SAMtools contained 244 individuals (DRX071278-DRX0771548) of an F2 population from an initial cross between *Oryza sativa* ssp. *japonica* cv. Nipponbare and the African wild rice species *O. longistaminata*, described by Furuta *et al*., (2017). For this dataset reads were trimmed to 65 bp length.

The second data set included 100 cassava genotypes (SRR1717936-SRR171973) from a bi-parental F1 population as described by the International Cassava Genetic Map Consortium (2015). All reads were trimmed to 85 bp after the cut-site.

Both datasets were processed using GBSmode described below. For comparison with SAMtools the data was trimmed in the same way as GBSmode and aligned using Bowtie2 (V 2.3.2, Langmead & Salzberg, 2012) with default settings. To make fair comparisons, the unfiltered output (the vcf files) were filtered using an R script that discarded loci with less than 50% of individuals genotyped and loci with less than five percent minor allele frequency.

### 5.2 Tag frequency

The GBSmode data analysis pipeline, shown in Fig. 1, takes as a starting point de-multiplexed fastq files (rawData directory) from a population. As GBS barcode size varies, GBSmode trims the reads to a uniform length and the frequency of each unique sequence is counted. This is done using the Bash command “grep –A1 ‘^@’¦ grep –v ‘-’¦ grep –v ‘@’¦ cut –c-85¦ sort¦ uniq –c”, where the first ‘grep’ command is a pattern match, which recognises the fastq header lines and returns them with their sequence data. The remaining ‘grep’ commands remove the headers and additional output. The ‘cut’ command selects the length of sequence to be extracted (85 bp in the example below). The sequences are then sorted alphabetically (using ‘sort’) before passing the output to the ‘uniq’ command, which counts (-c) the frequency of each unique sequence. Below is an example which will search a directory (rawData), containing the fastq data, and output the data to a new directory (inputDirectory).

~~~
for x in $(ls rawData/);
  do grep –A1 ‘^@’ rawData/$x | grep –v ‘-’ | grep –v
‘@’ | cut –c-85 | sort | uniq –c > inputDirectory/$x
done
~~~

**Figure 1.**
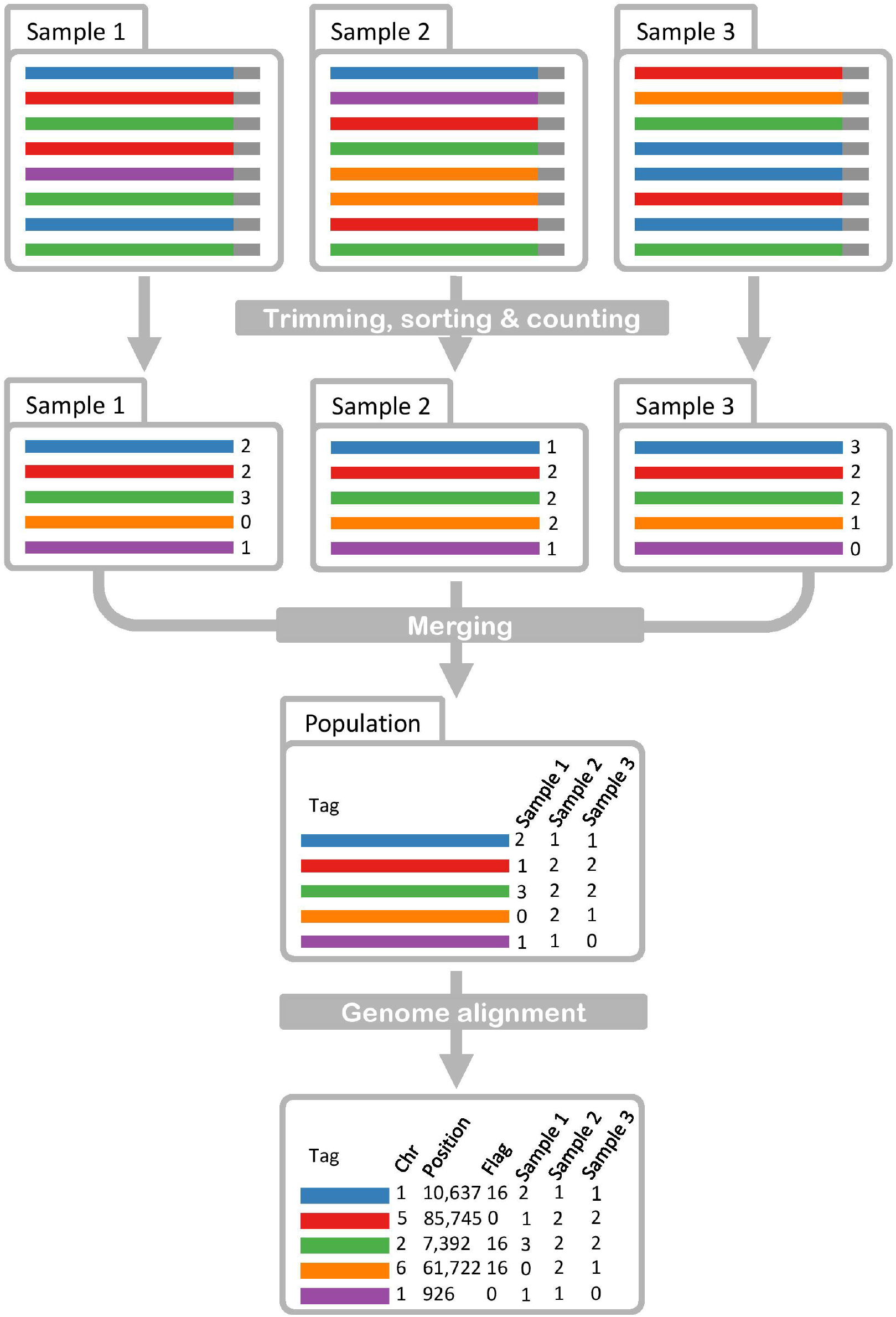
Schematic of the GBSmode pipeline. The schematic begins with raw data that is then trimmed, sorted and counted to produce a table of abundance of each unique read, per sample (Trimming, sorting & counting). As a next step, the data for each sample is combined into a table for the whole population (Merging). The reads are then aligned to the genome, to give coordinates for each read. This data includes the chromosome (Chr), position (bp) and flag, describing the read orientation (0=forward, 16= reverse).

### 5.3 Population counts

Next, the sample read counts are summarised per population, generating a matrix of unique sequences (rows) with genotype counts (columns) using the Perl script “getUniqueTags.pl”. This script exploits the hash data structure, in Perl, for speed and efficiency. A hash data structure contains unique keys with an associated value (like a dictionary). This Perl script uses a so-called ‘hash of hashes’ where the unique nucleotide sequences are the keys and within each key is another hash, with its key being the genotype identifier and its value the frequency of each count. In this form the data is stored and assigned easily before it is output, by iterating through the hash of hashes. When no information (counts) for a unique sequence is found for a genotype, it is reported as a zero.

~~~
perl getUniqueTags.pl inputDirectory/ > Count.data
~~~

### 5.4 Alignment

The data is back transformed into reads for aligning them to a reference. The Perl script “makeFastq.pl” converts the sequence data into fastq format, by assigning a unique name and gives a Phred quality score of 40 per base. The reads are then aligned to the genome using Bowtie2 (V 2.3.2, Langmead & Salzberg, 2012), producing a sequence alignment file (sam) with only the best alignment reported. The sam file is filtered for alignments to the genome (by selecting reads that map to chromosomes) and the sam file header is omitted, using ‘grep’ commands. This is to reduce computing time and memory requirements in the following R steps. The Perl script “TrimCountsInput.pl”, selects only reads that are mapped on the genome.

~~~
perl makeFastq.pl Count.data > sequences.fastq
bowtie2-build genome.fasta genome.ref
bowtie2 –x genome.ref –U sequences.fastq –S file.sam
grep “Chr” file.sam | grep –v “^@” | cut –f 1-6 | grep
‘Hap’ > filter.sam
perl TrimCountsInput.pl filter.sam Count.data >
filter.data
~~~

### 5.5 Biological factors

Genotyping is made using the “GBSmode.R” program, which can be run interactively in R or via a terminal. For example:

~~~
R –vanilla –slave “—args filter.data filter.sam fastq
genotype.delimiter genome.fasta” < GBSmode.R
~~~

with ‘fastq” being a common file identifier for the genotype names. Prior to genotype calling, in R, numerical factors are set to constrain the genotyping: the basal ploidy level of the genotypes (ploidy, 2), the maximum number of alleles allowed in the population (for example, two or four alleles in offspring of homozygous or heterozygous parents, respectively), minimum allele frequency (minAF, 100) and the minimum allele count (minAC, 8), which are described later.

### 5.6 Orientation

For genotyping, the alignment data (sam), population read-counts and genome sequence are required. The genome sequence is imported using ‘read.fasta’ from the “seqinr” package of R (Charif & Lobry, 2007). By default, the starting position of reads are left-most based in sam format, meaning the reads mapped in reverse orientation must be corrected. Therefore, the right-most position of reads (reverse) aligned on the negative strand corresponds to their origin-site position (Pos). This position is determined using the ‘extractAlignmentRangesOnReference’ function from the “GenomicAlignments” package (Lawrence *et al*., 2013), which uses the CIGAR (Concise Idiosyncratic Gapped Alignment Report) string to correct for insertions and deletions. To identify all unique positions, the direction (Flag), reference contig/chromosome (Ref), and origin-site position data are concatenated together to produce a unique position identifier (Upos). To eliminate low coverage sites only unique positions (Upos) with at least one tag greater than the minimum allele frequency (minAF, 100) are used.

### 5.7 Filtering

Each position (Upos) is validated based upon abundance of reads, maximum number alleles and prevalence among genotypes. Given that some reads may contain errors, only those which occur at a frequency greater than 5%, of all reads found at a Upos, are considered informative tags (Itags). A Upos is considered not polymorphic if the number of Itags is one. If a single genotype has more Itags than its ploidy (2 by default), or if the number of observed alleles (Itags) exceeds the maximum number of alleles possible (in the population), then the Upos is skipped.

### 5.8 Allele Alignments

To identify polymorphisms, the Itags are aligned together using the CIGAR string from the sam file using the ‘sequenceLayer’ function in “GenomicAlignments”. Any pairwise mismatches are considered polymorphic loci and termed informative sites (Isite). In some cases, the alignment renders Isites or reads redundant. Thus, the Upos again is checked; to be polymorphic, does not exceed the ploidy for individuals and that enough individuals can be genotyped.

### 5.9 Genotyping

For error correction, of reads, each unique read is assigned to a corresponding Itag, if it matches perfectly all Isite(s) positions from an Itag: otherwise it is removed. For genotyping, if a diploid genotype has two different Itags, then it is considered heterozygous. Individuals are homozygous when they have a single Itag with a frequency greater than 8 (minAC) (1-(^1^/_2_^(minAC-1)^) > 99% confidence).

The results are saved to two files, the genotyping and the count data (used for genotyping). To make the results comparable in this study and compatible with existing downstream analysis tools, we created the R script ‘ModeToVCF.R’ to convert the GBS mode output to variant call format (vcf).

### 5.10 Phasing

In the rice population, polymorphisms for individuals were phased by exploiting the population design. Where at any loci in the genome, of an individual, three possible states can exist; either it is heterozygous and contains both parental genomes, or homozygous for either the *O*.*japonica* or *O*.*longstaminata* parent: and should segregate in a 1:2:1 ratio. The phasing occurred in windows of 100 SNP loci, with an offset of 50 loci between consecutive windows. Within each window, a similarity matrix of the proportion of matches (ranging between 0-1) between genotypes was determined. The resulting similarity matrix was transformed into a (Euclidean) distance matrix. This distance matrix was organised using hierarchical agglomerative clustering (‘Ward’ method), before splitting into three groups (‘cutree’). Afterwards, the results were edited so the same cluster was consistent throughout a chromosome. This was made by permuting all cluster names in a proceeding window and selecting the ones that best match the prior. For consistency we manually edited for chromosomes so the heterozygous (most abundant) group was named the same way.

### 5.11 Comparison of variant calling

To compare the output of GBSmode and SAMtools, vcfR (v.1.12.0, Knaus & Grunwald, 2016) was used to load the data into R. To find exact matches, the chromosome, position, reference allele, and alternative allele fields were concatenated into one string. After removing the duplicates, the strings between GBSmode and SAMtools were compared to find one-to-one matches. To identify differences between how polymorphisms are reported, just the chromosome and position fields were concatenated together and compared. Closer examination of these results showed they were attributable to differences in how insertions and deletions are reported. For instance, SAMtools concatenates adjacent deletions together (e.g. “AAA”). Whereas GBSmode, when converted to vcf format, breaks adjacent deletions into separate polymorphisms (e.g. “A”, “A”, “A”). In addition, overlapping insertion and deletions, between datasets, were cross checked using their start and end point to find differences between reporting. Another type of difference occurred when filtering the vcf files for 50% of individuals being genotyped and greater than 5% minor allele frequency. To identify these errors, we loaded the un-filtered vcf files and repeated the same analysis as before when comparing filtered vcf files.

Finally, we re-ran GBSmode, but modified the script to output the reason for it rejecting loci. From these data we then looked for polymorphism found by SAMtools only and if they overlapped with a locus rejected by GBSmode (one hundred base pairs up- or downstream of the digestion site). From which we assigned them as being rejected for; ploidy reasons, because one or more sample contain three alleles: or it was found to be a singleton, meaning no variation was found (after filtering) by GBSmode.

## 6 Results

Using GBS data from an F2 population of rice (Furuta *et al*., 2017), the performance of GBSmode was benchmarked against SAMtools and identified, unfiltered, 15,187 polymorphic loci with a missing value rate of 51%. Of these loci, 8,554 contained a single SNP while 3,536, 1,603 and 705 contained two, three or four SNPs, respectively. For comparison, the GBSmode output was converted into vcf format and filtered for loci with at least 50 % of the individuals genotyped, yielding 13,449 polymorphic loci (Fig. 2a). Likewise, the same data was analysed using SAMtools (Li *et al*., 2009) and filtered using the same criteria. This yielded 14,445 polymorphic loci (Fig. 2a). In the work described by Furuta *et al*., (2017), TASSEL-GBS (Glaubitz *et al*., 2014) was used and identified 5,812 polymorphic loci with 50% missing value rate.

**Figure 2.**
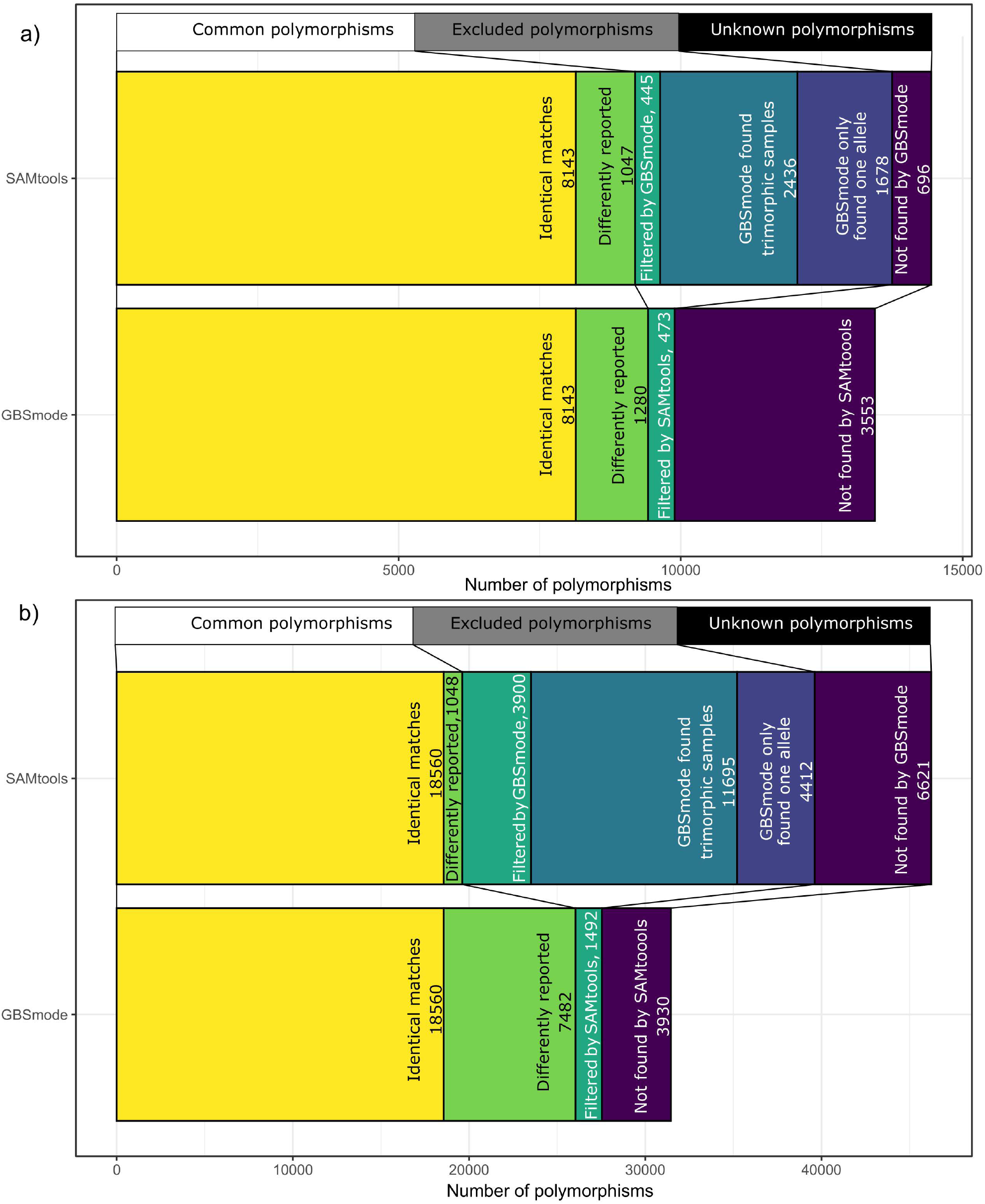
Stacked bar plots showing the overlap and differences between the output of GBSmode and SAMtools. The stacked bar plots show the cumulative number of polymorphic loci on the x-axis and the genotyping method on the y-axis. The stacks can be divided into three subdivisions shown above the bar plots. These include common polymorphisms, either one-to-one matches (shown in yellow) or those differently reported (green); excluded polymorphisms, those excluded when filtering the data for minimum number of genotypes (teal) and those excluded by GBSMode because they were found to be trimorphic (turquoise) or not polymorphic (blue); the final division is the number of unexplained loci (purple). Figure 3a shows the results using the rice data set with 244 individuals and figure 3b using 100 individuals using a cassava dataset.

To test the utility of the genotyping data, the overlapping genome segments were used to build phased haplotypes of the population, whereby each window, per genotype, was classified as heterozygous or homozygous for either the *O. japonica* or *O. longstaminata* parent. Using this approach, 264 and 274 haplotype windows were created by GBSmode and SAMtools, respectively (Fig. 3). The largest group of haplotypes (green) accounted for 29,862 haplotype windows for GBSmode and 32,564 for SAMtools. This was similar to the amount of the second and third largest haplotype-windows combined, with 25,770 for GBSmode and 28,436 for SAMtools. Thus, the data appear to fit the expected 1:2:1 segregation ratio. To verify the phasing, the frequency of consecutive blocks of three haplotype windows were counted. As expected, the consecutive haplotype blocks of three widows did not change 87% of the time. The haplotype windows switched between heterozygous and homozygous states in 8% of the cases. Likewise, in 4.5%, of the haplotype-windows the state switched from one homozygous state to the other, suggesting a double recombination in close proximity in both gametes or errors. The remaining 0.5% haplotype-window blocks switched between all three states or the middle haplotype window differed from the first and last. Whilst biologically possible it could be that these represent errors. Despite some discrepancies, it was evident the phasing had built a broad schematic of the genetic makeup of individuals.

**Figure 3.**
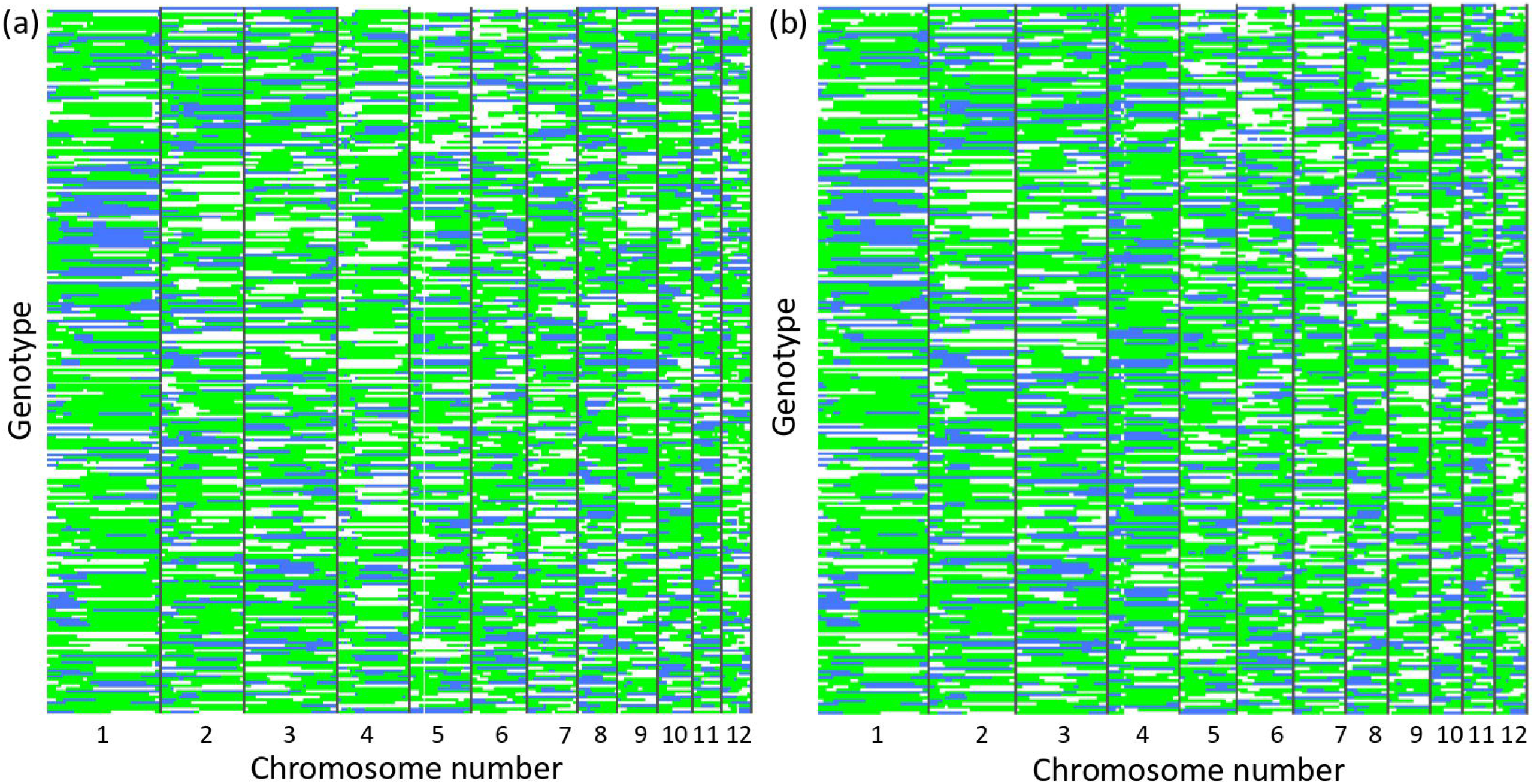
Results from phasing the rice population. The figure shows the results of phasing 244 individual rice genotypes (lines). Each block (column) represents the assignment of genotypes into three groups; heterozygous (green) or homozygous (white and blue). The phasing results from GBSmode is shown in panel a) and SAMtools in panel b).

The above phased data allowed us to determine the error rates of GBSmode and SAMtools. By intersecting the loci calls with expected haplotype-window data (using only consecutively matched haplotype-window blocks of three) the number of matches between the two was measured. For some loci the segregation pattern did not match that expected from the phasing, indicating reads misalignments. Therefore, the overlap between the haplotype-window and each locus was checked by Chi-squared test, using all individuals. In total 12,600 loci were tested for GBSmode and 13,700 for SAMtools. This found that 1,012 (8.0%) and 1,962 (14.3%) loci for GBSmode and SAMtools, respectively, did not show the same segregation pattern (FDR *P* < 0.05) as the phased haplotype-window. After removing these loci the mean number of correct calls was 95.0% and 91.7% for GBSmode and SAMtools, respectively. In another context, 2,642,778 genotyping calls were made for GBSmode, of which 211,422 are in wrong place and 120,638 are incorrect. SAMtools made 2,823,021 genotype calls, of which 404,290 were misplaced and 199,591 are incorrect. Thus, GBSmode does offer advantages for genotyping compared to SAMtools in terms of accuracy.

To follow up on the differences between GBSmode and SAMtools, the overlap between the called variants was assessed. After removing duplicates, the majority (8,143) of loci overlapped between GBSmode (13,449 total loci) and SAMtools (14,445 total loci). A further 1,280 and 1,047 were found in common between the datasets but reported differently in their vcf files. The number of polymorphisms absent in the filtered SAMtools vcf file but found in the GBSmode vcf file, which were subsequently found in the un-filtered SAMtools vcf file, was 473. Likewise, 445 polymorphims were found in the filtered SAMtools vcf file that were absent in the filtered GBSmode vcf file, but were present in the un-filtered GBSmode vcf file. In addition, we re-ran GBSmode and saved the reason for the GBSmode genotyping script excluding potentially polymorphic sites. The intersection of these data with the SAMtools vcf file found 2,436 polymorphisms were discarded by GBSmode because one or more individuals had three alleles, which is not biologically possible in a diploid sample. Finally, 1,165 polymorphisms were discarded by GBSmode, but found by SAMtools, because they were not polymorphic. These data suggest a substantial part (2,436/14,445) of the SAMtools data may be based upon read misalignments, which is similar to the number of loci which segregated differently (1,962).

To assess the utility of GBSmode in a more complicated biological system, a bi-parental F1 population of cassava, derived from two heterozygous diploid parents, was used. In data from 100 offspring, GBSmode identified 22,277 polymorphic loci, from which 2,706,052 genotype calls were made, with a 26.7 % missing-value-rate. When converted to vcf format (broken haplotypes) and filtering, 31,489 polymorphisms were reported, of which 176 were trimorphic and one tetramorphic. In comparison, SAMtools identified, after filtering, 43,680 polymorphic loci, of which 598 trimorphic and 52 tetramorphic markers. Further analysis of trimorphic and tetramorphic markers revealed that 315 of these markers were a single base in length. The remaining 267 contained more than 5 bp.

As with the rice dataset, the overlaps between the SAMtools and GBSmode results were compared (Fig. 3b). Many of polymorphisms (18,560) were exact one-to-one matches. Further investigation of common loci found 1,048 loci in the SAMtools dataset and 7,482 loci in the GBSmode dataset, that did overlap exactly but were reported differently. As described in the rice dataset, these are because of how insertions and deletions are reported. We then looked in the unfiltered vcf files to check for matches that had been excluded when filtering for minor allele frequency and minimum number of genotypes. Here, 3,900 loci were reported after filtering in the SAMtools dataset and absent in the filtered GBSmode dataset, but then found in the unfiltered GBSmode dataset. Likewise, 1,492 were found in the GBSmode filtered dataset which were only found in the SAMtools unfiltered dataset. This still meant 49% of polymorphisms (22,728/46,236) were unaccountable in the SAMtools dataset. To resolve this, we re-ran GBSmode and recorded the reason for the loci being excluded. The intersection of these data with the SAMtools dataset found most (51%, 11,695/22,728) had been discarded by GBSmode because individuals were found that contained three or more alleles (which is biologically impossible in a diploid individual). A further 4,412 alleles were discarded by GBSmode (but reported by SAMtools) because they were not polymorphic.

## 7 Discussion

Here we report a new method for the analysis of GBS data, called GBSmode. Using phased data from an F2 population of rice, we show evidence that GBSmode improves the genotyping accuracy by 9.6%. Because of the number reads incorrectly placed and incorrectly genotyped being reduced from 22.6% by SAMtools to 13.0% by GBSmode. Moreover, using an F1 cross in cassava, SAMtools discovered 43.7 K polymorphic loci, whereas GBSmode found 31.5 K polymorphic loci. But, 12.0 K of the loci reported by SAMtools were rejected by GBSmode as some samples contained three alleles (or more). Thus, genotyping by GBSmode yields higher quality by both removing erroneous read alignments and errors in genotyping.

One important finding of this work is the number of polymorphic loci where the segregation pattern of the SNP differed to that expected. In rice we found that 14% of loci were misplaced and in cassava up to 27%. This issue is attributable to both whole genome duplications and local duplications (Li *et al*., 2008). This must be kept in consideration when imputing SNPs, particularly by filling in missing data from those observed in the parents: which would propagate errors. Also, the number of miscalled genotypes was 5.0% and 8.3% for GBSmode and SAMtools respectively. In both cases, > 99% accuracy was set as a threshold. The difference between expected and observed error rates is attributable to low coverage sequencing (Scheben *et al*., 2017), whereby reads are only found from one parent in a heterozygous individual. For both cases (misplaced reads and genotyping errors) the best method to alleviate this is imputation and error correction. As shown here, imputation using blocks of reads is advantageous.

Throughout the development of this pipeline, a generally lower number of identified loci was observed compared to other genotyping software. As shown here, GBSmode is more stringent in calling genotypes, resulting in less polymorphic loci with less the errors. Whereas sometimes the trade-off between more errors outweighs the benefits more markers. For classic bi-parental populations or structured populations (such as a nested association mapping population), only a few thousand polymorphic loci are sufficient, especially when imputation is possible (Hyma *et al*., 2015; Furuta *et al*., 2017), so GBSmode would be more appropriate. In unstructured populations (namely genome-wide-association-studies) higher marker densities are needed to overcome linkage disequilibrium; thus, other tools might be preferable (Scheben *et al*., 2017). However, different genotyping tools are not exclusive and results from different genotyping pipelines can be combined. Whereby the vcf file generated by GBSmode can be integrated with other genotyping files produced by other pipelines.

A number of tools already exist for GBS analysis, for example SAMtools which is an all-purpose high throughput sequence analysis tool. Otherwise, TASSEL is designed for speed and for screening large populations. GBSmode leverages on accuracy and ease of use yet we foresee there are still margins of improvements for both aspects. First, summarising the population data can require significant amounts of random access memory (RAM) when the population data is tabulated. The step does not require much time, but genome complexity and sequencing errors can rapidly inflate the amount of RAM needed. In the future, the RAM requirements could be reduced by use of a SQL database. The second area of optimisation is the genotype calling, which runs sequentially in R. For the cassava dataset, this step took approximately two days. Parallelisation (multithreading in R) of loci screening could be a way to speed this up. A further improvement would be to make use of paired-end read data, currently reads are broken and analysed separately. The combination of forward and reverse reads would allow increased accuracy in read mapping and afford further combining of polymorphisms into haplotypes. Despite some room for improvement, GBSmode offers many attractive features such as counting unique reads, which significantly reduces the read-mapping step, as used in TASSEL. In addition, genotyping might be adapted for polyploid samples, tetraploids, or more complex combinations.

Taken together the results show that using all data from reads can significantly reduce the error rate in genotyping. The genotype calling in R makes the GBSmode script more accessible to other researchers for modification. Thus, it can be developed and adapted more broadly, making GBSmode a versatile tool for researchers and plant breeders alike.

## 8 Availability and requirements

Project name: GBSmode

Project home page: https://github.com/stevenandrewyates/GBSmode (DOI:10.5281/zenodo.4041275)

Operating system(s): Linux Programming language: Perl, R, Bash Other requirements: NA

License: MIT License

## 9 List of abbreviations

CIGAR, Concise Idiosyncratic Gapped Alignment Report; GBS, genotyping-by-sequencing; Isite, informative site; Itag, informative tag; MAF, minor allele frequency; minAC, minimum allele count; minAF, minimum allele frequency; Pos, origin-site position; RADseq, restriction site-associated DNA sequencing; RAM, random access memory; Ref, reference contig (chromosome); sam, sequence alignment (file); SNP, single nucleotide polymorphism; SQL, Structured Query Language; Upos, unique position; vcf, variant call format.

## 10 Declarations

### 10.1 Ethics approval and consent to participate

Not applicable

### 10.2 Consent for publication

Not applicable

### 10.3 Availability of data and material

https://github.com/stevenandrewyates/GBSmode

### 10.6 Authors’ contributions

Both authors devised the work, concept of GBSmode and wrote the manuscript. SY wrote the scripts.

## 10.7 Acknowledgements

The authors acknowledge Dr Dario Copetti, Dr. Roland Kölliker and Dr. Timothy Sykes for helpful comments on the manuscript and work.

## 10.8 Authors’ information (optional)

## References

Baird NA, Etter PD, Atwood TS, Currey MC, Shiver AL, Lewis ZA, Selker EU, Cresko WA, Johnson EA. Rapid SNP discovery and genetic mapping using sequenced RAD markers. PloS one. 2008;3(10):e3376.

Charif D, Lobry JR. SeqinR 1.0-2: a contributed package to the R project for statistical computing devoted to biological sequences retrieval and analysis. InStructural approaches to sequence evolution 2007 (pp. 207–232). Springer, Berlin, Heidelberg.

Ekblom R, Wolf JB. A field guide to whole □genome sequencing, assembly and annotation. Evolutionary applications. 2014;7(9):1026–42.

Elshire RJ, Glaubitz JC, Sun Q, Poland JA, Kawamoto K, Buckler ES, Mitchell SE. A robust, simple genotyping-by-sequencing (GBS) approach for high diversity species. PloS one. 2011;6(5):e19379.

Furuta T, Ashikari M, Jena KK, Doi K, Reuscher S. Adapting genotyping-by-sequencing for rice F2 populations. G3: Genes, Genomes, Genetics. 2017;7(3):881–93.

Glaubitz JC, Casstevens TM, Lu F, Harriman J, Elshire RJ, Sun Q, Buckler ES. TASSEL-GBS: a high capacity genotyping by sequencing analysis pipeline. PloS one. 2014;9(2):e90346.

Hyma KE, Barba P, Wang M, Londo JP, Acharya CB, Mitchell SE, Sun Q, Reisch B, Cadle-Davidson L. Heterozygous mapping strategy (HetMappS) for high resolution genotyping-by-sequencing markers: a case study in grapevine. PLoS One. 2015;10(8):e0134880.

International Cassava Genetic Map Consortium. High-resolution linkage map and chromosome-scale genome assembly for cassava (Manihot esculenta Crantz) from 10 populations. G3: Genes, Genomes, Genetics. 2015;5(1):133–44.

Knaus BJ, Grunwald NJ. VcfR: a package to manipulate and visualize VCF format data in R. bioRxiv. 2016; 041277. Publisher Full Text.

Langmead B, Salzberg SL. Fast gapped-read alignment with Bowtie 2. Nature methods. 2012;9(4):357.

Li H, Ruan J, Durbin R. Mapping short DNA sequencing reads and calling variants using mapping quality scores. Genome research. 2008;18(11):1851–8.

Li H, Handsaker B, Wysoker A, Fennell T, Ruan J, Homer N, Marth G, Abecasis G, Durbin R. The sequence alignment/map format and SAMtools. Bioinformatics. 2009;25(16):2078–9.

Lawrence M, Huber W, Pages H, Aboyoun P, Carlson M, Gentleman R, Morgan MT, Carey VJ. Software for computing and annotating genomic ranges. PLoS computational biology. 2013;9(8):e1003118.

Mortazavi A, Williams BA, McCue K, Schaeffer L, Wold B. Mapping and quantifying mammalian transcriptomes by RNA-Seq. Nature methods. 2008;5(7):621.

Ramu P, Esuma W, Kawuki R, Rabbi IY, Egesi C, Bredeson JV, Bart RS, Verma J, Buckler ES, Lu F. Cassava haplotype map highlights fixation of deleterious mutations during clonal propagation. Nature genetics. 2017;49(6):959.

Rasheed A, Hao Y, Xia X, Khan A, Xu Y, Varshney RK, He Z. Crop breeding chips and genotyping platforms: progress, challenges, and perspectives. Molecular plant. 2017;10(8):1047–64.

Scheben A, Batley J, Edwards D. Genotyping□by□sequencing approaches to characterize crop genomes: choosing the right tool for the right application. Plant biotechnology journal. 2017;15(2):149–61.

